# Insulin receptor substrate 4 deficiency mediates the insulin effect on the epithelial magnesium channel TRPM6 and causes hypomagnesemia

**DOI:** 10.1101/2022.10.01.510479

**Authors:** Jing Zhang, Sung Wan An, Sudha Neelam, Anuja Bhatta, Mingzhu Nie, Claudia Duran, Manjot Bal, Femke Latta, Jianghui Hou, Joseph J. Otto, Julia Kozlitina, Andrew Lemoff, Joost Hoenderop, Michel Baum, Matthias T Wolf

## Abstract

The kidney is the key regulator of magnesium (Mg^2+^) homeostasis in the human body. In the distal convoluted tubule (DCT), the apical epithelial magnesium (Mg^2+^) channel TRPM6, determines how much Mg^2+^ is excreted in the urine. To better understand the regulation of human renal Mg^2+^ absorption we identified novel, potential interaction partners of TRPM6 by pursuing a liquid chromatography – tandem mass spectrometry (LC-MS/MS) proteomics approach.

We found insulin receptor substrate 4 (IRS4) enriched with TRPM6 tagged to glutathione S-transferase (TRPM6-GST) but not GST control. Physical interaction between IRS4 and TRPM6 was confirmed by co-immunoprecipitation. Applying microdissection of mouse tubules, we detected *Irs4* mRNA expression mostly in the DCT and to a lower degree in the proximal tubule and thick ascending limb of Henle. Given the overall low abundance of *Irs4* mRNA along the tubule we investigated the phenotype of *Irs4* knockout mice (*Irs4*^-/-^). These *Irs4*^-/-^ mice displayed significantly higher urinary and fecal Mg^2+^ losses and lower blood Mg^2+^ levels than wild-type (WT) mice. Claudin-16, claudin-19, and Hnf1b mRNA and Claudin-16 and Trpm6 protein expression was significantly higher in kidneys of 3 month old *Irs4*^-/-^ mice consistent with a compensatory mechanism to conserve Mg^2+^. Applying whole-cell patch-clamp recording we confirmed the stimulatory role of insulin on TRPM6 channel activity and showed that IRS4 targets the two TRPM6 phosphorylation sites T1391 and S1583 to enhance TRPM6 current density. To test the effect of Mg^2+^ deficiency on metabolism, we performed glucose and insulin tolerance studies, which were mildly abnormal in *Irs4*^-/-^ mice.

**SIGNIFICANCE STATEMENT:** Magnesium (Mg^2+^) is the second most abundant intracellular cation but the regulation of Mg^2+^ homeostasis is not well understood. The kidney is the key organ for regulating Mg^2+^ homeostasis. Insulin is a known stimulator of the apical epithelial Mg^2+^channel TRPM6. We present a novel modifier of Mg^2+^ absorption with insulin receptor substrate 4 (IRS4) which illuminates further, how insulin activates the TRPM6 channel and modifies Mg^2+^ homeostasis. Applying protein biochemistry, tubular microdissection, whole mouse physiology, and patch-clamp recording, we demonstrate that IRS4 mediates the stimulatory effect of insulin by enhancing phosphorylation of two specific TRPM6 residues. *Irs4*^-/-^ mice develop increased urinary and stool Mg^2+^ losses, lower serum Mg^2+^ concentration, and display mild impairment in glucose and insulin tolerance.

## INTRODUCTION

Magnesium (Mg^2+^) represents the second most abundant intracellular cation in the human body but regulation of human Mg^2+^ homeostasis is not well understood (1, 2). Mild to moderate hypomagnesemia is frequently underappreciated because it does not display acute symptoms (3). Therefore, hypomagnesemia remains mostly undiagnosed and untreated, but occurs in up to 15% of the population (4, 5). Conditions associated with chronic hypomagnesemia include diabetes mellitus, metabolic syndrome, kidney disease, hypertension, and cancer (6–9). There is increasing evidence that hypomagnesemia correlates with type 2 diabetes mellitus (T2DM), gestational diabetes, and complications of diabetes such as diabetic nephropathy, retinopathy, foot ulcers, and coronary heart disease (7, 10–16). In the Atherosclerosis Risk in Community Study, hypomagnesemia doubled the odds for T2DM (10). Patients with T2DM have a 30% higher frequency of hypomagnesemia compared to the general public and increased urinary Mg^2+^ losses (7, 17, 18). However, the etiology of urinary Mg^2+^ losses in diabetes mellitus is unclear and may include poor Mg^2+^ intake, glomerular hyperfiltration, osmotic diuresis, and metabolic acidosis (19).

The gastrointestinal (GI) tract absorbs only about 50% of the oral Mg^2+^ intake of and loses the remaining 50% in the stool (1). The kidneys are more efficient in retaining Mg^2+^ and usually excrete only 3-5% of the filtered Mg^2+^ in the urine (1, 2). Therefore, the kidney is the major regulator of Mg^2+^ homeostasis. The majority (40-70%) of filtered Mg^2+^ is absorbed in the thick ascending limb (TAL) in a paracellular fashion driven by the lumen positive potential (1, 2). The fine-tuning of Mg^2+^ absorption occurs in the distal convoluted tubule (DCT) where about 5-10% of filtered Mg^2+^ is absorbed via the apical transient receptor potential melastatin 6 (TRPM6) and TRPM7 channel complex (20, 21). This is the final step of tubular Mg^2+^ absorption and determines the urinary Mg^2+^ excretion. Recessive mutations in *TRPM6* result in severe hypomagnesemia in infancy often causing seizures (22, 23). *Trpm6* knockout mice (*Trpm6*^-/-^) mice are embryonic lethal and develop neural tube defects (24).

Consistent with the association between hypomagnesemia and T2DM, human *TRPM6* single nucleotide polymorphisms (SNPs) are associated with T2DM and gestational diabetes mellitus (25, 26). These SNPs modify how well insulin can activate TRPM6 channel activity directly by phosphorylation (26). Other modifiers of TRPM6 regulation include hormones such as estrogen and insulin, which enhance TRPM6 mRNA transcription and modify TRPM6 phosphorylation, respectively (26, 27). The epidermal growth factor (EGF) affects TRPM6 mRNA expression and redistributes TRPM6 from endomembrane to plasma membranes (28, 29). ADP-ribosylation factor-like GTPase (ARL15) enhances TRPM6 channel activity and modifies metabolism (30). Medications such as thiazides, rapamycin, and calcineurin inhibitors reduce the TRPM6 mRNA abundance (31–34). *Ncc*-deficient mice, mimicking the Gitelman syndrome phenotype, are characterized by less TRPM6 mRNA and protein expression (34, 35). Recently, a new cause of a Gitelman-like phenotype was identified with mutations altering mitochondrial complex IV activity, which also affect TRPM6 (36). To better understand TRPM6 regulation we performed liquid chromatography - tandem mass spectrometry (LC-MS/MS) proteomic analysis of TRPM6-GST pulldown assays to identify new regulators of TRPM6.

## METHODS

### Materials and DNA constructs

Human TRPM6 was either transfected in a bicistronic vector pCINeo/IRES-GFP plasmid or in a pcDNA pDEST53 plasmid (ThermoFisher Scientific, Waltham, MA) (26, 37). Mutagenesis of TRPM6 in pCINeo/IRES-GFP was performed as published (26). Antibodies against human IRS4 (sc-100854) and TRPM6 were purchased from Santa Cruz Biotechnology (Santa Cruz, Dallas, TX). Antibody against Claudin-16 was kindly provided by Jianghui Hou (Washington University, St. Louis). For co-immunoprecipitation experiments a magnetic beads Dynabeads Immunoprecipiation kit (ThermoFisher Scientific, Waltham, MA) was used. Mouse monoclonal anti-β-actin-peroxidase antibody were purchased from Sigma (Saint Louis, MO). For knockdown experiments of IRS4 siRNA was used from Dharmacon (Lafayette, CO). We purchased the mitogen-activated protein kinase (MAPK) and PKB inhibitor ML-9 from Enzo Life Sciences (Farmingdale, NY).

### Cell culture and transfection, siRNA-mediated knockdown of IRS4, and insulin treatment

HEK293 cells were cultured as described (37). Cells were transiently transfected using Lipofectamine 2000® reagent (Thermo Fisher Scientific, Waltham, MA) with plasmids (1-2 μg per 6-well) containing TRPM6-GFP, or control vectors as indicated. The total amount of transfected DNA was balanced by using empty vectors. To study the effect of IRS4 HEK293 cells were transfected with non-targeting siRNA or 10 nM siRNA targeting human IRS4 (Dharmacon, Lafayette, CO) using Lipofectamine 2000 per manufacture protocol. Experiments were performed 48 h after transfection. To assess the effect of insulin on TRPM6 current density, cell medium was changed 32 h after transfection to serum-free Opti-MEM. The next day cells were incubated for 1 h with 30 nM insulin (Millipore-Sigma, Saint Louis, MO) prior to whole-cell patch-clamp recording.

### LC-MS/MS analysis of potential TRPM6 interaction partners

The human TRPM6 cDNA was cloned into the pGEX-KG plasmid (ATCC, Manassas, VA) providing an N-terminal GST-tag. We separated cell lysates from HEK293 cells overexpressing empty N-terminal GST or human N-terminal GST-TRPM6 on a SDS-PAGE gel. After silver-staining a 10 mm gel slice, the appropriate protein band was cut and diced. Protein gel pieces were reduced and alkylated with DTT (20 mM) and iodoacetamide (27.5 mM) as described (38). A 0.1 μg/μl solution of trypsin (Thermo Fisher Scientific, Waltham, MA) in 50 mM triethylammonium bicarbonate (TEAB) was added to cover the gel, allowed to sit on ice, and then 50 μl of 50 mM TEAB was added and the gel pieces were digested overnight at 37°C. An Oasis HLB μelution plate (Waters, Milford, MA) was used for solid-phase extraction cleanup. Resulting peptides were reconstituted in 10 μl of 2% (v/v) acetonitrile (ACN) and 0.1% (v/v) trifluoroacetic acid in water. Two microliters of this were injected onto an Orbitrap Fusion Lumos spectrometer (Thermo Electron, West Palm Beach, FL) coupled to an Ultimate 3000 RSLC-Nano liquid chromatography system (Dionex, Sunnyvale, CA). Samples were injected onto a 75 μm i.d., 75 cm long EasySpray column (Thermo Fisher Scientific, Waltham, MA), and eluted with a gradient from 1-28% buffer B over 90 minutes. Buffer A contained 2% (v/v) ACN and 0.1% (v/v) formic acid in water, and buffer B contained 80% (v/v) ACN, 10% (v/v) trifluoroethanol, and 0.1% (v/v) formic acid in water. The mass spectrometer operated in positive ion mode. MS scans were acquired at 120,000 resolution in the Orbitrap, and up to 10 MS/MS spectra were obtained in the ion trap for each full spectrum acquired using higher-energy collisional dissociation for ions with charges 2-7. Dynamic exclusion was set for 25 s after an ion was selected for fragmentation. Raw MS data were analyzed using Proteome Discoverer v2.4 (Thermo Fisher Scientific, Waltham, MA), with peptide identification performed using Sequest HT searching against the human protein database from UniProt. Precursor and fragment tolerances of 10 ppm and 0.6 Da were specified and three missed cleavages were allowed. Carbamidomethylation of Cys was set as a fixed modification, with oxidation of Met set as a variable modification. The false-discovery rate cutoff was 1% for all peptides. Protein abundances were determined by summing the peak intensities for all peptides matched to each specific protein and were used to compare relative protein amounts between samples.

### Co-immunoprecipitation and immunoblotting

To allow for TRPM6 to form protein complexes with endogenous IRS4, HEK293 cells were transfected with TRPM6 or control plasmid using Lipofectamine 2000 and incubated for 48 h at 37°C. After transfection, the cells were lysed with 500 μl of RIPA lysis buffer on ice and resuspended in 450 μl of IP extraction buffer (Thermo Fisher Scientific, Waltham, MA) with 100 mM sodium chloride, 2 mM magnesium chloride, 1 mM DTT, and 5 μl of protease inhibitors. The cells were incubated on ice for 5 min and centrifuged at 2500 g for 5 min at 4°C. Supernatant was transferred to a fresh tube for Co-IP. The protein concentration in the cell lysate was determined using protein assay (DC Protein Assay, Biorad, Hercules, CA). Samples were adjusted to the same concentration with buffer. For immunoprecipitations, we used a magnetic beads Dynabeads Immunoprecipiation kit (Thermo Fisher Scientific, Waltham, MA). Dynabeads were coupled with antibody using the antibody coupling protocol. Briefly, 5 mg of Dynabeads were washed in the C1 buffer (Thermo Fisher Scientific, Waltham, MA) by vortexing and placed on the magnet column for 1 minute. After settlement of the beads supernatant was discarded. Then either 5 μg of anti-IRS4 or anti-TRPM6 antibody (Santa Cruz, Dallas, TX) was added to the beads together with C2 buffer to a final volume of 500 μl. The tube was incubated on a shaker at 37°C overnight. The next day the beads were washed twice and resuspended in 100 μl of storage buffer (Thermo Fisher Scientific, Waltham, MA). We transferred 1.5 mg of (TRPM6 or IRS4) antibody–coupled beads to a tube and added 20 μg of cell lysate. The mixture was incubated on a shaker at 4 °C for 30 min. The tubes were placed on a magnetic column for 1 min. Once the beads settled supernatant was discarded. Dynabeads were washed 3× in 200 μl of extraction buffer. After the final wash, 60 μl of elution buffer was added to the tubes. Tubes were incubated on a shaker for 5 min at room temperature and then placed on the magnetic column for 1 min to allow settlement of the beads. The supernatant containing the purified protein complex was then resolved on 4-20% gradient gel and processed for immunoblotting using specific antibodies. Species-specific anti-IgG (Santa Cruz, Dallas TX) was used as control. Samples were denatured at 65°C for 10 minutes (min) and subjected to SDS-PAGE and immunoblotting. Antibodies against human TRPM6 (Santa Cruz, Dallas, TX) and IRS4 (Santa Cruz, Dallas, TX) were used for Western blotting.

### Animals

*Irs4*^-/-^ mice were purchased from the Jackson Laboratory (Bar Harbor, ME). These mice were published by the Lienhard laboratory (39, 40). Because the mouse *Irs4* gene is localized on the X chromosome we focused on studying male animals as we anticipated weaker phenotypes in female *Irs4*^-/-^ mice. Male mice were genotyped using tail clippings and were studied at 3, 6, and 12 months. All animal experiments were performed in adherence to the NIH Guide for the Care and Use of Laboratory Animals and were approved by the University of Texas Southwestern Medical Center at Dallas Institutional Animal Care and Use Committee.

### Microdissection studies

Kidneys from 4-6 week-old *Irs4*^-/-^ and WT mice were removed after mice were anesthetized with intraperitoneal Inactin (10 mg/100 g body weight). Removed kidneys were cut into several coronal slices for microdissection. Slices of kidney were placed into a prewarmed collagenase type I (Worthington, Lakewood, NJ; 1.5 mg/ml dissolved in DMEM/F12) solution in a 15-ml tube, which was then shaken vigorously on a titer plate shaker at 37°C for 10-15 minutes. After digestion, the individual nephron segments were dissected in ice-cold Hanks solution containing (in mM) 137 NaCl, 5 KCl, 0.8 MgSO_4_, 0.33 Na_2_HPO_4_, 0.44 KH_2_PO_4_, 1 MgCl_2_, 10 Tris (hydroxymethyl) amino methane hydrochloride, 0.25 CaCl_2_, 2 glutamine, and 2 L-lactate (pH 7.4) and transferred by adhering the tubules to small glass beads (0.5 mm diameter, Thomas Scientific, Swedesboro, NJ) (41, 42). Beads were then transferred to 1.5 ml tubes containing 0.6 ml RNase inhibitor-containing lysis buffer. Proximal tubules, thick ascending limb of Henle, and distal convoluted tubules were dissected free hand in Hank’s solution. Their RNA extraction was performed using Quick-RNA microPrep (Zymo Research, Irvine, CA). Reverse transcription and qRT-PCR were performed as outlined further below. Sequences of primers for RT-PCR analysis were published (37, 43).

### Metabolic cage studies in mice

Blood and urine electrolytes were determined in *Irs4*^-/-^ and WT mice at three, six, and twelve-month on a C57BL/6 background (39). Animals were acclimated for three days in metabolic cages (Tecniplast, West Chester, PA) prior to experiments. Mice were handled and monitored as described (37). Urine electrolytes except Mg^2+^ were analyzed by the Metabolic Phenotyping Core of the UTSW Touchstone Diabetes Center using a Vitros®350 Chemistry System Clinical Analyzer. Urine Mg^2+^ and creatinine were determined by the UTSW O’Brien Kidney Center applying atomic absorption and capillary electrophoresis, respectively. Urine volume was normalized for urinary creatinine.

### Phlebotomy and blood analysis

Blood samples were obtained from mice anesthetized with isoflurane. Via retroorbital bleed one hundred microliters was collected in heparinized tubes for analysis of electrolytes. Serum was tested for electrolytes including urine by the Metabolic Phenotyping Core of the UTSW Touchstone Diabetes Center using a Vitros®350 Chemistry System Clinical Analyzer. Serum creatinine was analyzed by the UTSW O’Brien Kidney Center applying capillary electrophoresis.

### Quantitative RT-PCR and RT studies

For total kidney RNA, tissue was isolated from kidneys from WT and *Irs4*^-/-^ mice using miRNeasy Mini kits from Qiagen (Germantown, MD). First-strand cDNA was synthesized by iScript™ cDNA synthesis kit (Bio-Rad, Hercules, CA). In control reactions, we ensured that no amplification was produced when reverse transcriptase was replaced by water. Relative transcript expression was measured by quantitative real-time PCR using iTaq™ Universal SYBR® Green Supermix (Bio-Rad, Hercules, CA). Samples were run on CFX96 Real-Time PCR Detection System (Bio-Rad, Hercules, CA). 18S RNA was used to normalize for expression of mRNA. Primers for qRT-PCR were published (37, 43). Data were analyzed using the Bio-Rad CXF software and calculated by ΔΔCt value. For the nested RT-PCR approach we synthesized cDNA as described above. Aliquots of the RT and control reactions were amplified by PCR using published mouse Irs4 primers (Irs4 F: TTGCCGCCTCAGAGCTATCC and Irs4 R: GTGCTGAGCGTCGGAAAAGG), 2.5 U AmpliTaqGold (Perkin Elmer) in a 20 μl volume. Amplification consisted of 3 min at 95°C, 35 cycles of 10 sec at 95°C, 30 sec at 55°C, and elongation with a gradient ranging from 65°C to 95°C with an increment of 0.5 °C for 1 min, with a final 7 min at 72°C. An aliquot of 5μl of each PCR reaction was reamplified for 30 cycles using the same protocol as for the first amplification. 10 μl of the final reaction mix was separated on a 1.5% (w/v) agarose gel and visualized with ethidium bromide applying a Gel Doc XR+ Gel Documentation system (Bio-Rad).

### Whole-cell patch-clamp recording

In brief, whole-cell patch-clamp recordings were performed in a similar fashion as outlined (44, 45). Approximately 48 h after transfection HEK293 cells were dissociated and placed in a chamber for ruptured whole-cell recordings as described. Transfected cells were identified for recording by their GFP fluorescence. Similar to previous publications we used TRPM6 bath solution containing (in mM) 140 NaCl, 5 CsCl, 1 MgCl_2_, 10 glucose, 10 HEPES (pH 7.4 with NaOH), and the pipette TRPM6 solution containing (in mM) 140 NaCl, 10 EDTA, and 10 HEPES, (pH 7.2 with NaOH) (20, 46, 47). Whole-cell patch-clamp pipettes were pulled from borosilicate glass (Sutter Instrument, Novato, CA) and had resistance between 1.5-3 MΩ. The cell membrane capacitance and series resistance were monitored and compensated (>75%) electronically using an Axopatch 200B amplifier (Axon Instruments, Foster City, CA). Voltage protocol consists of 0 mV holding potential and successive voltage sets (200-ms duration) from −100 to +100 mV in +20 increments. Current densities were obtained by normalizing current amplitude (obtained at +100 mV) to cell capacitance. Data acquisition was performed using ClampX10.4 software (Axon Instruments). Currents were low-pass filtered at 2 kHz using 8-pole Bessels filter in the clamp amplifier, sampled every 0.1 ms (10kHz) with Digidata-1440 interface, and stored directly to a computer hard drive. For stimulation by insulin cells were treated for h with 30 nM insulin.

### Statistical analysis

Student’s t-tests were used to test if there are significant differences in the continuous outcomes between two study groups. For multiple comparisons one-way ANOVA studies followed by Student-Newman-Keuls method, allowing for pairwise multiple comparisons, were performed. Data are reported as means ± S.D if not outlined differently. *P*<0.05 was considered statistically significant.

## RESULTS

### IRS4 interacts physically with the Mg^2+^ channel TRPM6

In two LC-MS/MS experiments, we identified IRS4 as a potential GST-TRPM6 interaction partner (Fig. 1A). IRS4 was also an intriguing candidate for TRPM6 regulation given the regulation of TRPM6 by insulin (26). To confirm the physical interaction between TRPM6 and IRS4 we performed co-immunoprecipitation experiments of the TRPM6-IRS4 protein complex (Fig. 1B). The TRPM6 protein formed a protein complex with endogenous IRS4. We used anti-IRS4 antibody to immunoprecipitate IRS4 and co-immunoprecipitated TRPM6 (upper panel of Fig. 1B). In the reverse experiment we immunoprecipitated TRPM6 with anti-TRPM6 antibody and co-immunoprecipitated IRS4 (lower panel of Fig. 1B). These experiments confirmed physical interaction between TRPM6 and IRS4 as suggested by LC-MS/MS.

**Figure 1.**
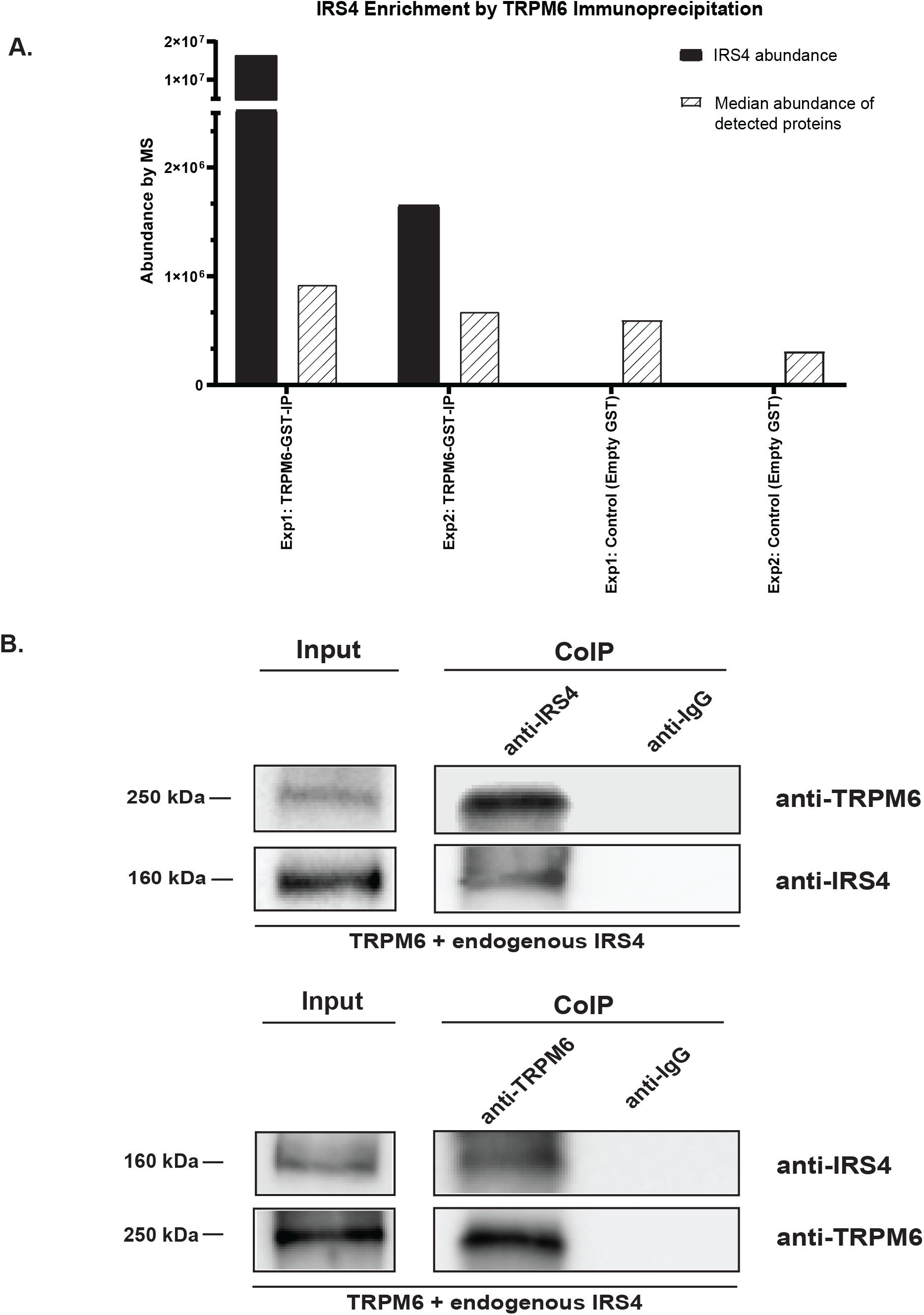
IRS4 and TRPM6 physically interact. (A) In two separate LC-MS/MS experiments (Exp1 and Exp2) we identified IRS4 enriched with GST-TRPM6 compared to the median abundance of detected proteins. We did not identify any IRS4 in HEK293 cells transfected with empty GST plasmid as control. In the first set of experiments (Exp1 in Fig. 1A), IRS4 abundance was 1.65×10^7^ compared to a median abundance of other detected proteins of 9.28×10^5^. In a second experiment (Exp2 in Fig. 1A) IRS4 abundance was 1.65×10^6^ compared to a median abundance of detected proteins of 6.79×10^5^ (Fig. 1A). The median abundance of detected proteins with empty GST plasmid were 6.02×10^5^ and 3.14×10^5^, with no detection of IRS4. (B) We confirmed the potential physical interaction between endogenous IRS4 and transfected TRPM6 by co-immunoprecipitation. HEK293 cells were transfected with pcDNA pDEST53 TRPM6 plasmid. When precipitating the TRPM6-IRS4 complex with anti-IRS4 we also detected TRPM6 at approximately 250 kDa. Performing the reverse experiment, we precipitated the TRPM6-IRS4 complex with anti-TRPM6 and identified IRS4 at approximately 160 kDa.

### IRS4 mRNA is expressed the mouse DCT

As we identified IRS4 as an interaction partner of TRPM6, we hypothesized that *Irs4* mRNA is localized in the mouse DCT, the nephron segment where *Trpm6* mRNA expression is most abundant (22). We studied relative *Irs4* mRNA abundance in the PT, TAL, and DCT using qRT-PCR and nested RT PCR (Fig. 2). Identification of individual tubules was based on morphology and relative location. Nephron site was confirmed by analysis of mRNA abundance of segment-specific markers (*Nkcc2* for TAL and *Ncc* for DCT) (Fig. 2A). As expected *Nkcc2* mRNA abundance was highest in the TAL and *Ncc* mRNA abundance was maximum in the DCT. Consistent with our hypothesis, we detected the most *Irs4* abundance in the mouse DCT with no detection of *Irs4* in the PT and TAL due to exceeding the numbers of amplification cycles with qRT-PCR. Due to this limitation, we further investigated potential low degree *Irs4* mRNA expression in the PT or TAL with a nested RT-PCR approach (Fig. 2B). In post-fetal mouse tissue, it is well known, that *Irs4* mRNA expression is not very abundant and requires a nested amplification approach (39). Performing reverse transcription followed by two rounds of amplification and normalizing for housekeeping gene *18S* expression we detected weaker *Irs4* bands in the mouse PT and TAL with 28% and 30% compared to DCT, respectively (Fig. 2B). The highest *Irs4* mRNA abundance was again in the DCT confirming the strong *Irs4* mRNA expression in the DCT from the qRT-PCR approach (22).

**Figure 2.**
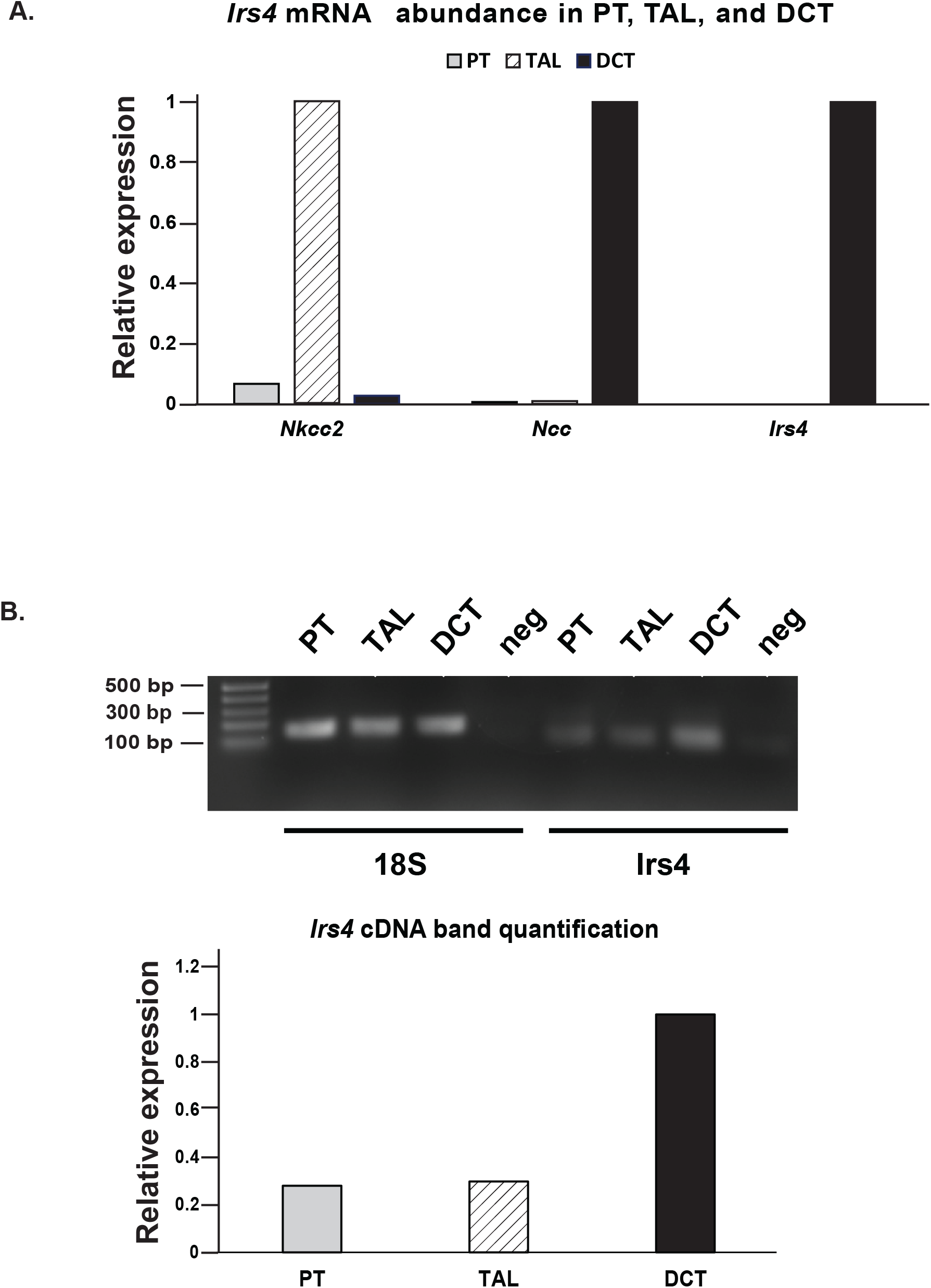
Irs4 mRNA is localized in the mouse DCT. (A) Microdissected tubules from WT mice displayed appropriate mRNA expression of tubule-specific markers NKCC2 for the TAL and NCC for the DCT. Applying qRT-PCR, we found the highest Irs4 mRNA abundance in the mouse DCT, the same nephron segment where Trpm6 mRNA is highly expressed. (B) As the low Irs4 mRNA tissue expression limited the qRT-PCR approach with no Irs4 detection in the PT and TAL, we performed a nested RT-PCR approach. Correcting the tubule-specific Irs4 cDNA expression for the housekeeping gene expression 18S we confirmed the highest Irs4 abundance in the DCT with lower Irs4 expression in the mouse PT (28%) and TAL (30%) compared to DCT. The negative (neg) control showed no significant PCR product after the nested amplification. *n*=10 tubules for each.

### Irs4^-/-^ mice are characterized by urinary and fecal Mg^2+^ losses and develop hypomagnesemia

Given the low abundance of *Irs4* in post-fetal tissue, we wondered if the lack of *Irs4* had any physiological significance in whole mouse physiology (39). Therefore, we examined urinary Mg^2+^ excretion (Fig. 3A), serum Mg^2+^ (Fig. 3B), and fecal Mg^2+^ (Fig. 3C) excretion in *Irs4*^-/-^ and WT mice at 3, 6, and 12 months. Similar to humans, the *Irs4* gene is on the murine X chromosome and so a milder phenotype is anticipated in human and murine females. While male *Irs4*^-/-^ mice have a significantly lower weight than WT mice, this was not found in female *Irs4*^-/-^ mice (40). Therefore, we focused only on studying *Irs4*^-/-^ male mice. At 3 months, we detected a significantly higher urinary excretion of Mg^2+^ in *Irs4*^-/-^ mice compared to WT animals. However, at this age, there was no significant difference regarding serum Mg^2+^ (Fig. 3B) or fecal Mg^2+^ excretion (Fig. 3C). At 6 months, we identified a significantly higher urinary Mg^2+^ excretion in *Irs4*^-/-^ mice compared to WT animals. This was accompanied by a significantly lower serum Mg^2+^ in *Irs4*^-/-^ mice compared to WT mice and higher fecal Mg^2+^ losses. At 12 months, we detected again a significantly higher urinary excretion of Mg^2+^ in *Irs4*^-/-^ mice compared to WT animals. This was accompanied by a significantly lower serum Mg^2+^ in *Irs4*^-/-^ mice compared to WT mice and higher fecal Mg^2+^ losses. A summary of all serum and urine electrolytes is provided in Table 1. Similar to Fantin el al., we noticed a slightly but significantly lower weight in 3 month-old *Irs4*^-/-^ compared to WT mice which persisted at 6 and 12 months (40). Serum K^+^ was lower at 6 and 12 months but there were no changes in urinary K^+^excretion at 6 months. Moreover, we also noticed significantly higher serum glucose values at 6 and 12 months (Table 1).

**Figure 3.**
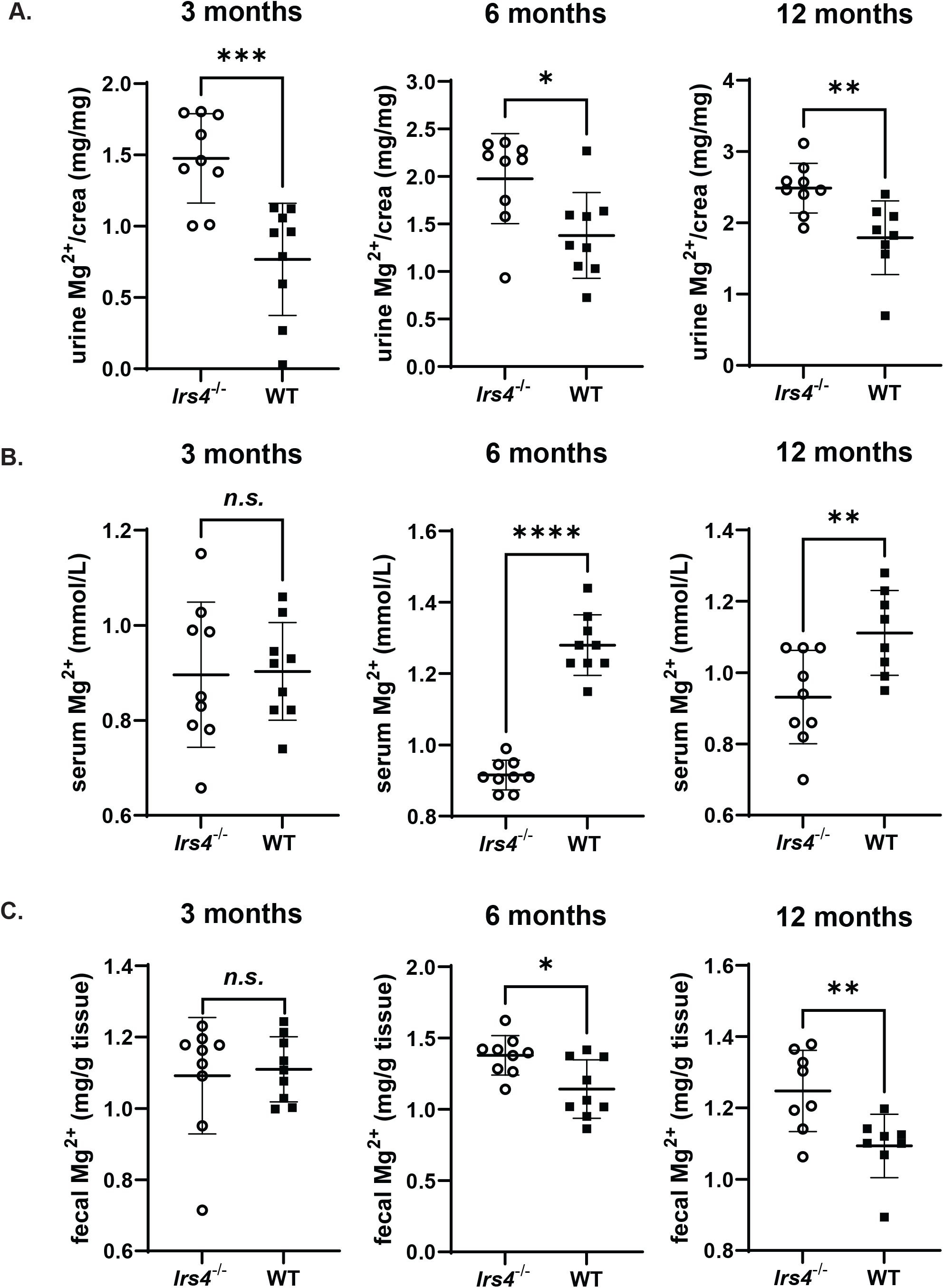
*Irs4*^-/-^ mice display urinary and fecal Mg^2+^ losses and develop lower serum Mg^2+^ levels. (A) Urine Mg^2+^ corrected for urinary creatinine was significantly higher at 3 (1.48±0.31 vs. 0.77±0.39 mg/mg for *Irs4*^-/-^ vs. WT, *p*<0.001), 6 (1.98±0.47 vs. 1.38±0.45 mg/mg for *Irs4*^-/-^ vs. WT, *p*<0.05), and 12 months (2.49±0.35 vs. 1.79±0.52 mg/mg for *Irs4*^-/-^ vs. WT, *p*<0.005) in male *Irs4*^-/-^ compared to WT mice. (B) At 3 months, serum Mg^2+^levels were not significantly different between *Irs4*^-/-^ and WT mice (0.90±0.15 vs. 0.90±0.15 mmol/L for *Irs4*^-/-^ vs. WT, *n.s*.). However, at 6 (0.92±0.04 vs. 1.28±0.08 mmol/L for *Irs4*^-/-^ vs. WT, *p*<0.0001) and 12 months (0.93±0.13 vs. 1.11±0.12 mmol/L for *Irs4*^-/-^vs. WT, *p*<0.01) serum Mg^2+^ values were significantly lower in *Irs4*^-/-^ compared to WT mice. (C) At 3 months, no significant difference was found in fecal Mg^2+^ excretion (1.09±0.16 vs. 1.11±0.09 mg/g feces for *Irs4*^-/-^ vs. WT, *n.s*.). At 6 (1.38±0.14 vs. 1.14±0.2 mg/g feces for *Irs4*^-/-^ vs. WT, *p*<0.05) and 12 months (1.25±0.11 vs. 1.09±0.09 mg/g tissue for *Irs4*^-/-^ vs. WT, *p*<0.01) *Irs4*^-/-^ mice displayed a higher fecal excretion of Mg^2+^ compared to WT mice. *n*=8-9 per group. * *p*<0.05; ** *p*<0.005; *** *p*<0.001; **** *p*<0.0001

**Table 1.** Summary of weight, urine, and blood data from Irs4-/- and WT mice at 3, 6, and 12 months.

|  | 3 months |  | 6 months |  | 12 months |  |
| --- | --- | --- | --- | --- | --- | --- |
|  | *Irs4*<sup>-/-</sup> | Wild-type | *Irs4*<sup>-/-</sup> | Wild-type | *Irs4*<sup>-/-</sup> | Wild-type |
| Weight (gram) | 26.3 ± 1.5* | 27.9 ± 1.3 | 30.2 ± 3.3<sup>\$</sup> | 35.2 ± 2.4 | 31.5* ± 3.2 | 35.9 ± 3.4 |
| **Urine excretion** |  |  |  |  |  |  |
| Mg<sup>2+</sup>/creatinine (mg/mg) | 1.48 ± 0.31<sup>#</sup> | 0.77 ± 41.4 | 1.98 ± 0.47* | 1.38 ± 0.45 | 2.49 ± 0.35<sup>\$</sup> | 1.79 ± 0.52 |
| FE Mg<sup>2+</sup> (%) | 0.08 ± 0.02* | 0.05 ± 0.01 | 0.08 ± 0.01<sup>\$</sup> | 0.05 ± 0.02 | 0.13 ± 0.02<sup>\$</sup> | 0.07 ± 0.02 |
| Na<sup>+</sup>/creatinine (mg/mg) | 16.8 ± 2.6 | 15.7 ± 2.9 | 11.5 ± 2.9* | 14.7 ± 2.7 | n.d. | n.d. |
| K<sup>+</sup>/creatinine (mg/mg) | 49.2 ± 7.6 | 47.5 ± 6.7 | 45.3 ± 8.9 | 46.9 ± 6.4 | n.d. | n.d. |
| Ca<sup>2+</sup>/creatinine (mg/mg) | 0.34 ± 0.17 | 0.26 ± 0.14 | 0.28 ± 0.13 | 0.24 ± 0.07 | n.d. | n.d. |
| **Serum** |  |  |  |  |  |  |
| Na<sup>+</sup> (mmol/L) | 152 ± 5 | 155 ± 4 | 151 ± 2 | 150 ± 1 | 153 ± 3<sup>\$</sup> | 150 ± 1 |
| K<sup>+</sup> (mmol/L) | 5.1 ± 0.4 | 4.9 ± 0.4 | 4.8 ± 0.4* | 5.2 ± 0.3 | 4.3 ± 0.6* | 4.9 ± 0.5 |
| Ca<sup>2+</sup> (mmol/L) | 2.48 ± 0.06 | 2.44 ± 0.06 | 2.36 ± 0.04* | 2.4 ± 0.04 | 2.4 ± 0.06 | 2.43 ± 0.09 |
| Mg<sup>2+</sup> (mmol/L) | 0.9 ± 0.15 | 0.9 ± 0.1 | 0.92 ± 0.04<sup>#</sup> | 1.28 ± 0.08 | 0.93 ± 0.13* | 1.11 ± 0.12 |
| PO<sub>4</sub><sup>3-</sup> (mmol/L) | n.d. | n.d. | 1.79 ± 0.27* | 2.02 ± 0.15 | 1.80 ± 0.17 | 1.71 ± 0.32 |
| Glucose (mmol/L) | 11.6 ± 2.4 | 12 ± 2.2 | 9.8 ± 1.6<sup>#</sup> | 4.8 ± 1 | 9 ± 1.8* | 6.7 ± 1.2 |
| BUN (mmol/L) | n.d. | n.d. | 8.3 ± 1.3 | 7.4 ± 0.8 | 21.2 ± 4.8 | 20.7 ± 5.3 |
| Creatinine (μmol/L) | 5.8 ± 0.9 | 5.9 ± 0.8 | 5.4 ± 0.7* | 7.5 ± 1.7 | 7.5 ± 1.1 | 6.5 ± 1 |
| **Stool** |  |  |  |  |  |  |
| Mg<sup>2+</sup> (mg/g tissue) | 1.09 ± 0.16 | 1.11 ± 0.09 | 1.38 ± 0.14* | 1.14 ± 0.20 | 1.25 ± 0.11* | 1.09 ± 0.09 |

Mean with S.D., n.d., no data, n=8-9 mice; FE, fractional excretion
\* $p < 0.05$ , $^{\$}$ $p < 0.005$ , $^{\#}$ $p < 0.0005$

### Magnesiotropic genes are upregulated in Irs4^-/-^ mice

Given the serum and urine abnormalities of Mg^2+^ in *Irs4*^-/-^ mice, we tested if genes involved in tubular Mg^2+^ absorption are upregulated in *Irs4*^-/-^ mice to compensate for the urinary Mg^2+^ losses. Therefore, we examined mRNA and protein expression of different magnesiotropic genes/proteins in kidney tissue from male 3-month-old *Irs4*^-/-^ and WT mice as published (37). We identified significant upregulation of *Hnf1b, Claudin-16*, and *Claudin-19* mRNA abundance in kidneys of *Irs4*^-/-^ mice (Fig. 4A). No significant difference in mRNA abundance was found for *Fxyd2b, Egf, Parvalbumin,* and *Trpm6.* Trpm6 and Claudin-16 protein abundance was also significantly higher in *Irs4*^-/-^ mice (Fig. 4B and supplementary Fig. 1A & 1B). The upregulation of *Hnf1b, Claudin-16,* and *Claudin-19*mRNA and Trpm6 and Claudin-16 protein expression in *Irs4*^-/-^ mice is consistent with the attempt of the tubule to compensate for ongoing urinary Mg^2+^ losses.

**Figure 4.**
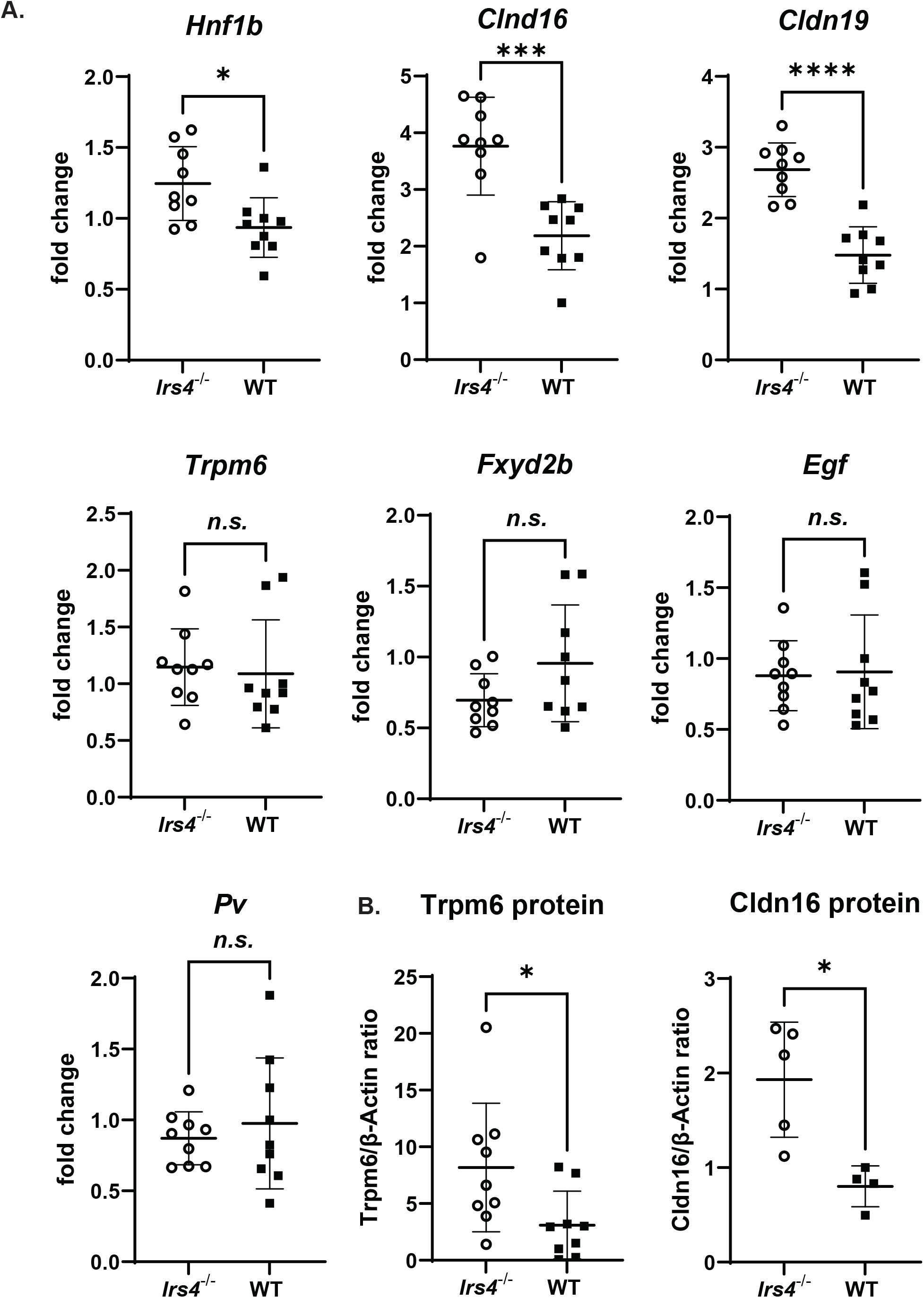
Magnesiotropic genes are upregulated in the kidneys of *Irs4*^-/-^ mice. (A) To test if the urinary Mg^2+^ losses in *Irs4*^-/-^ mice enhance transcription of magnesiotropic genes, we obtained kidneys from 3 months old male *Irs4*^-/-^ and WT mice and performed qRT-PCR. We detected significantly upregulated mRNA gene expression for Hnf1b, Claudin-16, and Claudin-19. No significantly different gene expression was detected for Fxyd2b, Egf, Parvalbumin, and Trpm6. (B) In total kidney lysates from 3 months old male *Irs4*^-/-^ and WT mice we identified a significantly higher protein expression for Trpm6 and Claudin-16 in *Irs4*^-/-^ compared to WT mice. *n*=4-9 per group. * *p*<0.05; ** *p*<0.005; *** *p*<0.001; **** *p*<0.0001

### IRS4 mediates the insulin stimulation of TRPM6

Next, we investigated the mechanism of how IRS4 modifies TRPM6. Our first goal was to confirm the stimulatory role of insulin on TRPM6. We studied whole-cell current density in HEK293 cells transfected with TRPM6. Insulin significantly enhanced TRPM6 whole-cell current density (Fig. 5A). To confirm that the TRPM6 stimulation was dependent on a signaling pathway relevant to insulin signaling we applied ML-9, a myosin light chain inhibitor, which blocks mitogen-activated protein kinase (MAPK) and PI3K-AKT/PKB (48, 49). When adding ML-9 to control and insulin-treated groups there was no significant stimulation of TRPM6 whole-cell current density (Fig. 5A). These findings indicate that TRPM6 stimulation by insulin occurs through a physiological meaningful, insulin-dependent pathway. Next, we tested if the insulin-mediated stimulation of TRPM6 depends on IRS4. We approached this question by applying siRNA knockdown of endogenous IRS4 in HEK293 cells. The efficiency of IRS4 siRNA knockdown was 97% compared to control siRNA (inlet of Fig. 5B). When cells were co-transfected with control siRNA and stimulated with insulin there was an increase in TRPM6 current density detected (Fig. 5B). However, when cells were co-transfected with IRS4 siRNA, insulin had no significant stimulatory effect on TRPM6 (Fig. 5B). These findings suggested that IRS4 mediates the stimulatory effect of insulin on TRPM6.

**Figure 5.**
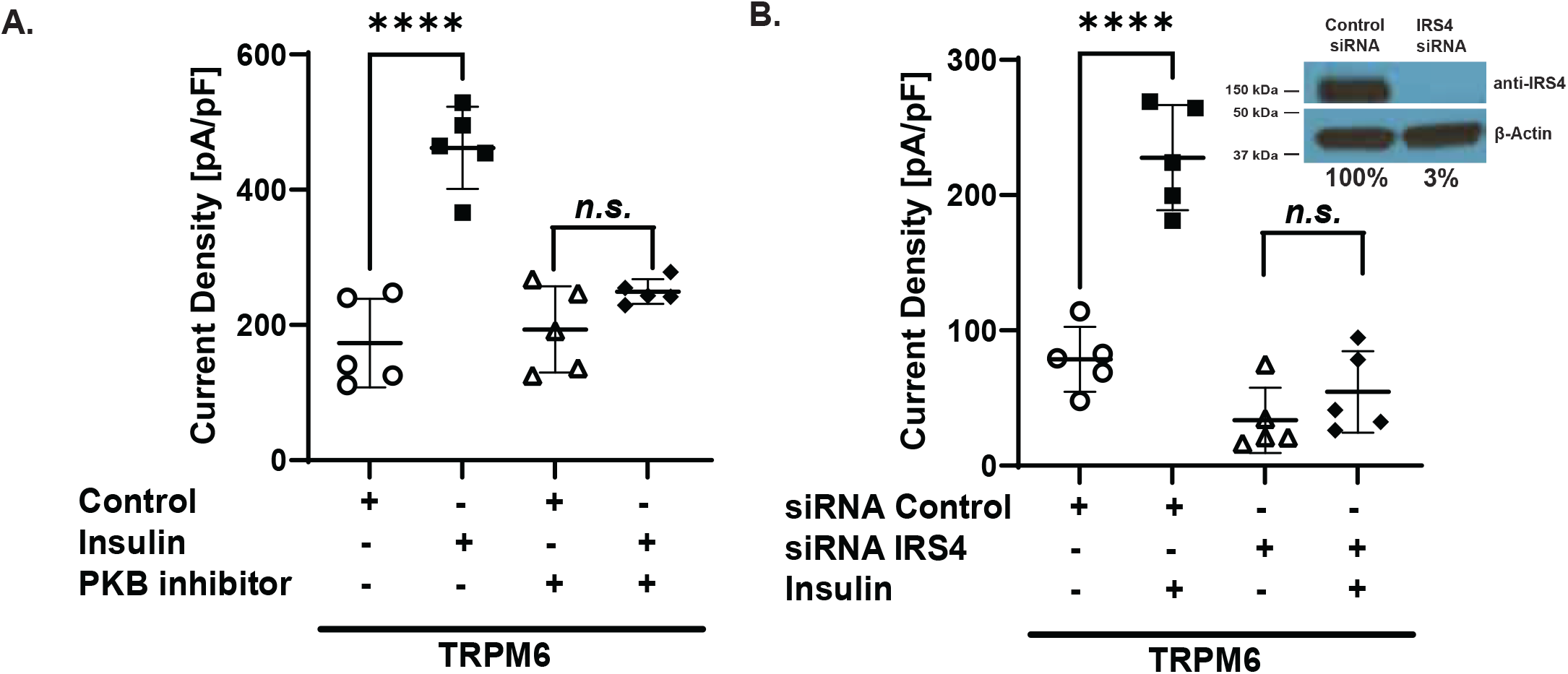
Insulin stimulates TRPM6 current density via IRS4. (A) Insulin significantly increased TRPM6 current density (173±66 vs. 461±61 pA/pF, control vs. insulin, *p*<0.0001) (26). When blocking the insulin signaling pathway with the PI3K-AKT/PKB inhibitor ML-9 stimulation of TRPM6 by insulin was abrogated (193±64 vs. 249±18 pA/pF, control vs. insulin, *n.s*.). (B) To test if the TRPM6 stimulation by insulin is mediated by IRS4 we knocked-down endogenous IRS4 in HEK293 cells by applying siRNA (inlet). Using control siRNA in TRPM6 transfected HEK293 cells insulin still enhanced TRPM6 current density (79±24 vs. 228±39 pA/pF, control siRNA with control vs. insulin, *p*<0.0001). When co-transfecting IRS4 siRNA insulin did not significantly enhance TRPM6 current density (34±24 vs. 55±30 pA/pF, control siRNA with control vs. insulin, *n.s*.). *n*=5 per group. **** *p*<0.0001

### IRS4 mediates phosphorylation of the TRPM6 residues T^1391^ and S^1583^

Nair et al identified two specific TRPM6 residues (T^1391^ and S^1583^), which are phosphorylated by insulin (26). These two residues are close to two SNPs V^1393^I and K^1584^E, which associate with gestational diabetes mellitus and abolish TRPM6 stimulation by insulin. These two SNPs were also found to correlate with a higher susceptibility for T2DM in a cohort of Korean women with low Mg^2+^ intake (25). To study if IRS4 is specifically involved in the insulin-mediated phosphorylation of the T^1391^ and S^1583^ residues we used either phosphorylation-deficient mutants (T^1391^A and S^1583^A) or variants mimicking the phosphorylated state (T^1391^D and S^1583^D). When co-transfected with control siRNA the phosphorylation-deficient TRPM6 mutants T^1391^A and S^1583^A were both not significantly activated by insulin (Fig. 6A & 6C). When applying IRS4 siRNA also no significant TRPM6 current stimulation was noticed for both mutants with insulin (Fig. 6A & 6C). We repeated these experiments for the phospho-mimetic TRPM6 variants T^1391^D and S^1583^D (Fig. 6B & 6D). When co-transfected with control siRNA the phospho-mimetic TRPM6 mutants T^1391^D and S^1583^D were both significantly activated by insulin (Fig. 6B & 6D). However, when applying IRS4 siRNA no significant TRPM6 current stimulation was noticed for both mutants after insulin treatment (Fig. 6B & 6D). Our patch-clamp experiments are consistent with a role for IRS4 in the phosphorylation of the two TRPM6 T^1391^and S^1583^ sites by insulin and point to IRS4 as a crucial downstream mediator of the insulin effect on TRPM6.

**Figure 6.**
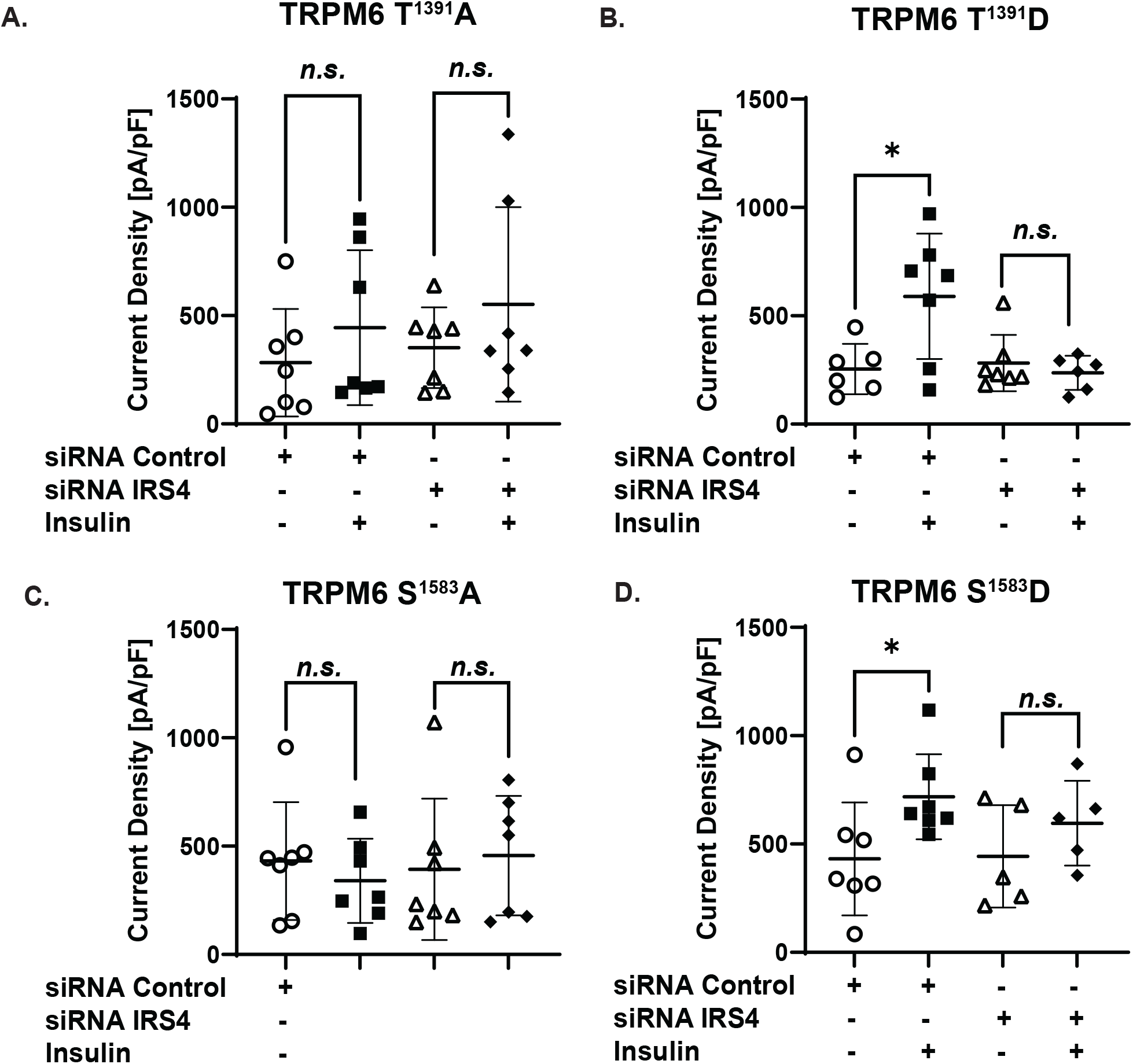
Insulin stimulates TRPM6 current density via phosphorylation of the TRPM6 residues T^1391^ and S^1583^. We evaluated the effect of IRS4 on the stimulation of phospho-deficient (T^1391^A and S^1583^A) and phospho-mimetic (T^1391^D and S^1583^D) TRPM6 residues. (A) Using the phospho-deficient TRPM6 variant T^1391^A no significant response was detected with insulin treatment and either with control siRNA or IRS4 siRNA (for T^1391^A: 283±249 vs. 445±357 pA/pF, control siRNA with control vs. insulin, *n.s*., and 352±186 vs. 552±448 pA/pF, IRS4 siRNA with control vs. insulin, *n.s*.). (B) When applying the phospho-mimetic TRPM6 variant T^1391^D stimulation with insulin increased TRPM6 current density significantly with control siRNA (for T^1391^D: 254±116 vs. 590±290 pA/pF, control siRNA with control vs. insulin, *p<0.05)* but not with IRS4 siRNA (for T^1391^D: 282±130 vs. 237±78 pA/pF, IRS4 siRNA with control vs. insulin, *n.s*.). (C) Similarly, transfecting the phospho-deficient TRPM6 variant T^1583^A no significant response was identified with insulin treatment and either with control siRNA or IRS4 siRNA (for S^1583^A: 432±272vs. 340±195 pA/pF, control siRNA with control vs. insulin, *n.s*., and 393±326 vs. 456±275 pA/pF, IRS4 siRNA with control vs. insulin, *n.s*.). (D) However, when cotransfecting the phospho-mimetic TRPM6 variant T^1583^D and control siRNA insulin enhanced TRPM6 current density (for S^1583^D: 431±261 vs. 718±196 pA/pF, control siRNA with control vs. insulin, *p*<0.05) while co-transfection with IRS4 siRNA did not increase TRPM6 current density significantly (for S^1583^D: 443±237 vs. 596±196 pA/pF, control siRNA with control vs. insulin, *n.s*.). *n*=5-7 per group. * *p*<0.05

### Deficiency of IRS4 is not compensated by higher IRS1-3 mRNA expression

In humans, there are six members of the IRS family (50). However, IRS5 and IRS6 are poor substrates for the insulin receptor (IR) and seem not to be relevant for insulin signaling (51, 52). Hence, we focused on IRS1-3. IRS1 has a major role in growth and IRS2 is required for insulin responsiveness of liver and muscle tissue. Both, IRS1 and IRS2, are more widely expressed. IRS3 is limited to the adipocytes and brain, and IRS4 is mostly distributed in embryonic tissues and cell lines (50, 53). IRS3 and IRS4 modify the actions of IRS1 and IRS2 by antagonizing some of their actions when expressed at high levels (54). Because the IRS1-4 proteins are homologous and characterized by multiple similar tyrosine-phosphorylation motifs we investigated if the lack of *Irs4*^-/-^ is compensated by upregulation of the other *Irs1-3* mRNAs. Therefore, we performed RT-qPCR of kidneys from WT and *Irs4*^-/-^ mice. We found no significantly different mRNA expression of *Irs1, Irs2,* or *Irs3* in the kidneys of WT versus *Irs4*^-/-^ mice (Fig. 7).

**Figure 7.**
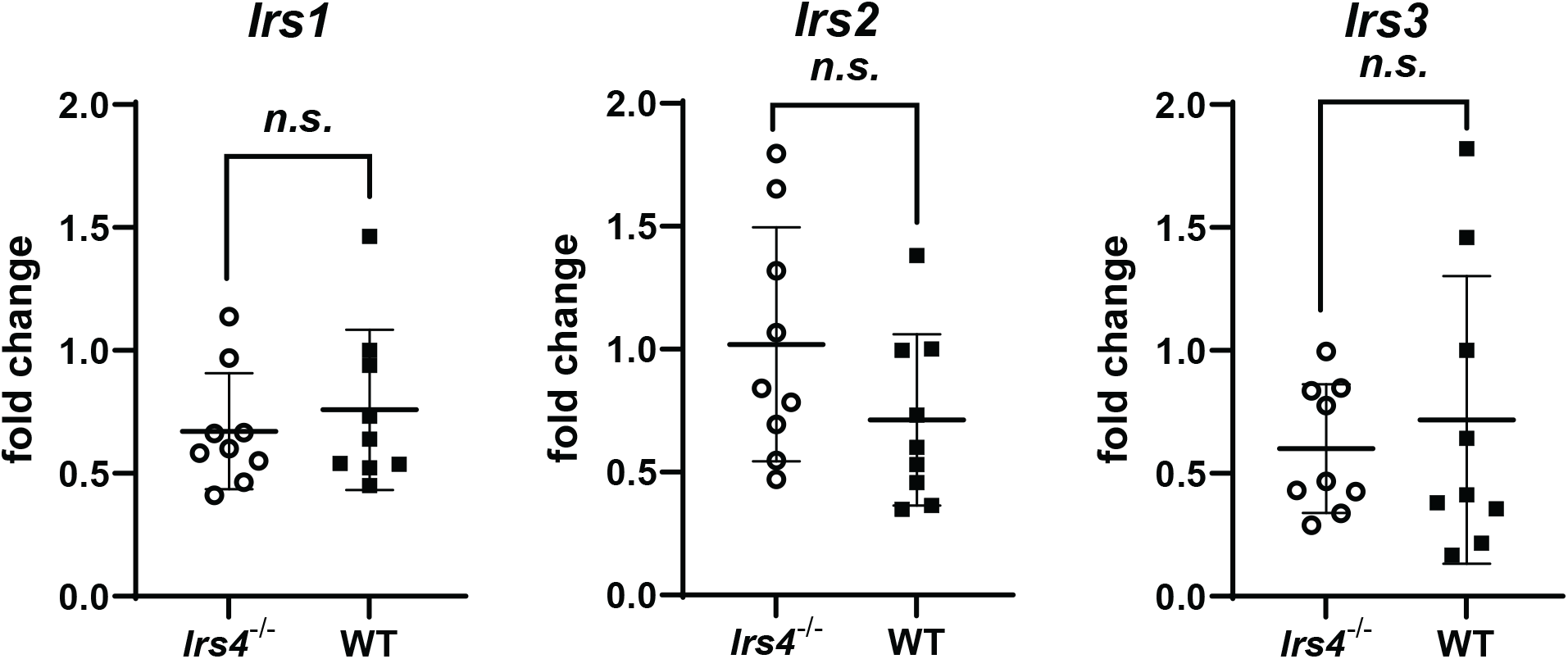
Gene expression of *Irs1, Irs2*, and *Irs3* is not significantly altered in the kidneys of *Irs4*^-/-^ mice. We evaluated kidneys from 3 month old male *Irs4*^-/-^ and WT mice for *Irs1, Irs2,* and *Irs3* mRNA expression. No significantly different mRNA expression for *Irs1*, *Irs2*, and *Irs3* was found in WT vs. *Irs4*^-/-^ mice. *n*=9 per group

### Glucose tolerance in Irs4^-/-^ mice is only modestly altered

To test if the Mg^2+^ depletion at 6 months impairs glucose tolerance, we performed a glucose tolerance test (GTT) (Fig. 8A). At 6 months, *Irs4*^-/-^ mice displayed a mildly impaired glucose tolerance at 30 and 120 min. To test if this was due to a lower sensitivity to insulin we performed an insulin tolerance test (ITT), which showed slightly better response to insulin in male *Irs4*^-/-^ mice at 30 and 120 min but overall no lack of response to insulin in *Irs4*^-/-^ mice when compared WT mice after insulin treatment (Fig. 8B).

**Figure 8.**
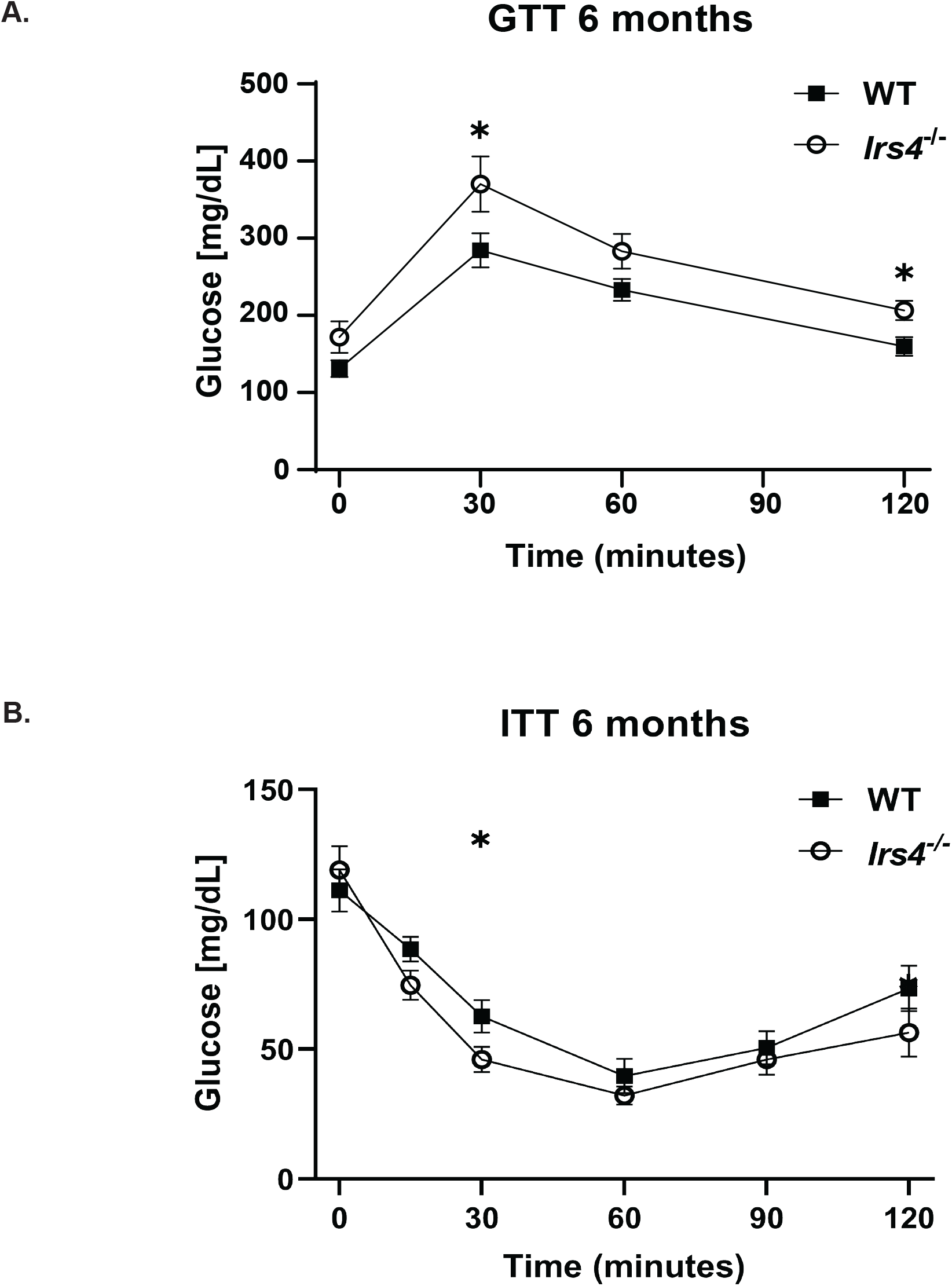
Glucose metabolism in *Irs4*^-/-^ mice. (A) We performed a glucose tolerance test after an overnight fast in male *Irs4*^-/-^ and WT mice at the age of 6 months and detected mildly impaired glucose tolerance. Data is shown as means ± S.E.M. *n*=8-9 per group. (B) Subsequently, we conducted an insulin tolerance test which showed no significant difference in male *Irs4*^-/-^ and WT mice in response to insulin. Data is shown as means ± S.E.M. *n*=8-9 per group. * *p*<0.05

## DISCUSSION

We describe the identification of IRS4 as a downstream member of the insulin-signaling cascade to regulate TRPM6. Nair et al. showed that the insulin receptor (IR) localizes to the DCT, the same nephron segment where TRPM6 is most abundant (26). Furthermore, two variants of the human *TRPM6* gene associate with T2DM and gestational diabetes (25, 26). IRS4 is primarily detected in embryonic tissue and primary cell lines (39, 53, 55).

Insulin affects electrolyte and water absorption by stimulating basolateral IRs in the proximal tubule, the DCT, and likely other nephron segments (56–60). Insulin signaling is initiated with insulin binding to the IR, a receptor tyrosine kinase, which then phosphorylates IRS proteins. IRS proteins are large scaffolding proteins which are recruited to the IR (50). IRS proteins mediate their action through stimulation of the PI3K-AKT/PKB or the MAPK pathways (19, 61). It has been hypothesized that one of the other IRS proteins in mice may compensate for the lack of IRS4 (40). Our data do not show any evidence for such a mechanism (Fig. 7). These findings are in line with data from different knockout mouse studies outlining that each IRS isoform has different biological consequences, supporting the idea that IRS1-4 function rather complementary than redundant in murine insulin signaling (62, 63). IRS4 is thought to counteract IRS1 and IRS2 actions, but similar to IRS1, IRS4 also translocates GLUT4 to the cell membrane and mediates PI3K-dependent metabolic actions of insulin (54, 64, 65).

*Irs4*^-/-^ mice display a surprisingly consistent phenotype regarding urinary and fecal Mg^2+^ losses and hypomagnesemia (Fig. 3), which was unexpected given the relatively low abundance of *Irs4* in post-embryonic tissue (Fig. 2) (39, 40). One could have expected a certain degree of compensation of Mg^2+^ homeostasis in 6 or 12 month-old *Irs4*^-/-^ mice.

However, given the fecal loss of Mg^2+^ in *Irs4*^-/-^ mice we think these mice are unable to compensate for the continued renal Mg^2+^ loss and subsequently develop hypomagnesemia at 6 and 12 month (Fig. 3B & 3C). It remains unclear why 6 and 12 month-old *Irs4*^-/-^ mice have fecal Mg^2+^ losses but 3 month-old *Irs4*^-/-^ mice do not. Fecal Mg^2+^ losses may be explained with a role of *Irs4* in the GI tract where Trpm6 is localized in the caecum and colon (27, 66, 67).

In humans, X-chromosomal mutations in *IRS4* result in central hypothyroidism with a milder and later phenotype in females (68). There is no data published on Mg^2+^ or glucose homeostasis in humans with *IRS4* mutations. However, considering IRS4 as the terminal part of insulin signaling it is intriguing that recessive loss-of-function mutations in the gene encoding the IR *(INSR)* cause Donohue and Rabson-Mendenhall syndromes which are characterized by insulin resistance, growth retardation, and low to borderline normal serum Mg^2+^ values – reminiscent of the *Irs4*^-/-^ mice (69). While patients with central hypothyroidism due to *IRS4* mutations have most likely loss-of-function mutations, human SNPs in *IRS4* have been associated with myogenesis and insulin-dependent tumor growth, which are possibly due to gain-of-function *IRS4* variants (70–73). This may also provide an explanation why the *Irs4* mRNA level is relatively low in post-fetal tissue and requires nested PCR amplification because higher *Irs4* expression may possibly contribute to a higher risk of tumorgenesis in adult mice.

The association between hypomagnesemia and diabetes mellitus has been known since the 1940s (74). However, the molecular mechanism how insulin and Mg^2+^ interact remains mostly unclear. In erythrocytes, insulin enhances Mg^2+^ absorption and cytosolic Mg^2+^ concentration (75). In cardiomyocytes and pancreatic β-cells, insulin stimulates glucose and Mg^2+^ uptake, suggesting a link between glucose and Mg^2+^ homeostasis (76, 77). Microperfusion experiments in the TAL and DCT showed increased Mg^2+^ absorption after insulin stimulation (58, 78). On the other hand, Mg^2+^ also modifies glucose metabolism. Mg^2+^ plays an important role in the autophosphorylation of the IR, as the receptor tyrosine kinase of the IR requires Mg^2+^ ions for enhancing the IR’s affinity to adenosine triphosphate (79–81). This was confirmed in hypomagnesemic rats, which had reduced levels of IR phosphorylation, resulting in insulin resistance (82, 83). Other Mg^2+^deficiency-related effects on glucose homeostasis include reduced glucose absorption via glucose transporter 4 (GLUT4), impaired insulin secretion, and altered lipid profiles (84–90). In rats, a low Mg^2+^/high fat diet caused Mg^2+^ disturbances, reduced protein phosphorylation of proteins in the insulin signaling pathway, and insulin resistance (91). In rats with streptozotocin-induced diabetes mellitus 16 weeks of MgSO4 therapy improved glucose tolerance, blood glucose, Akt2 and Irs1 gene and protein expression (92).

A role for Mg^2+^ in glucose metabolism is also suggested by patients with Gitelman syndrome, who have hypomagnesemia and a higher prevalence for T2DM (93). In addition, after solid organ transplantation, the use of calcineurin inhibitors, which cause renal Mg^2+^ wasting and hypomagnesemia, is also associated with a prevalence of new onset of diabetes mellitus after a solid organ transplant (NODAT) (33, 94–97). A higher susceptibility for metabolic syndrome and insulin resistance was also described in a large human cohort with higher urinary Mg^2+^ losses due to variant in *ARL15.* The encoded protein ARL15 enhances TRPM6 channel activity (30). *Irs4*^-/-^ mice display mild defects in growth, fertility, and glucose homeostasis (40). Our GTT and ITT results are comparable to published data in *Irs4*^-/-^ mice (40). Slightly lower serum glucose values were described 6-10 week old *Irs4*^-/-^ animals but only for the male mice the difference was significant (40). No significant differences for serum insulin were found in male and female *Irs4*^-/-^vs. WT animals at 6-10 weeks and 6 months. In a glucose tolerance test both male and female animals showed a slightly impaired glucose tolerance which were significant only for male but not for female animals (40). Future experiments in *Irs4*^-/-^ mice using low and high Mg^2+^ diets will further delineate the role of Mg^2+^ deficiency and therapy regarding glucose homeostasis.

## AUTHOR CONTRIBUTIONS

M.T. Wolf conceived and planned the study. J. Zhang, S.W. An, J. Hoenderop, J. Hou, A. Lemoff and M.T. Wolf designed the research. J. Zhang, S.W. An, S. Neelam, M. Nie, A. Bhatta, C. Duran, M. Bal, M. Baum, F. Latta, J. Otto performed the experiments. J. Zhang, S.W. An, S. Neelam, M. Nie, C. Duran, A. Bhatta, M. Bal, F. Latta, and J. Otto analyzed the data. A. Lemoff and J.J. Otto performed the LC-MS/MS analysis. S.W. An, J. Hoenderop, A. Lemoff, M. Baum, J. Kozlitina, and M.T. Wolf interpreted the results of the experiments. J. Zhang, S.W. An, J. Otto, and M.T. Wolf prepared the figures. M. Baum and M.T. Wolf drafted the manuscript. J. Zhang, S.W. An, S. Neelam, M. Nie, C. Duran, A. Bhatta, M. Bal, F. Latta, J. Otto, J. Hou, J. Hoenderop, A. Lemoff, J. Kozlitina, M. Baum, and M.T Wolf edited, revised and approved the final version of the manuscript.

## ACKNOWLEDGEMENTS

We thank Jiang Hui for providing us with the Claudin-16 antibody and Rose Nguyen for her technical support.

## DISCLOSURES

None

## FUNDING

Matthias T Wolf is supported by the Department of Defense (W81XWH1910205), the National Institute of Health (R01DK119631, DK079328-11), and the Children’s Clinical Research Advisory Committee (CCRAC), Children’s Health System, Dallas.

## SUPPLEMENTAL MATERIAL

See supplemental Figure 1 attached.

## Notes

### Competing Interest Statement

The authors have declared no competing interest.

