## Supplementary figures and images for "Insulin receptor substrate 4 deficiency mediates the insulin effect on the epithelial magnesium channel TRPM6 and causes hypomagnesemia"

### Supplemental Figure 1

Supplementary Figure 1

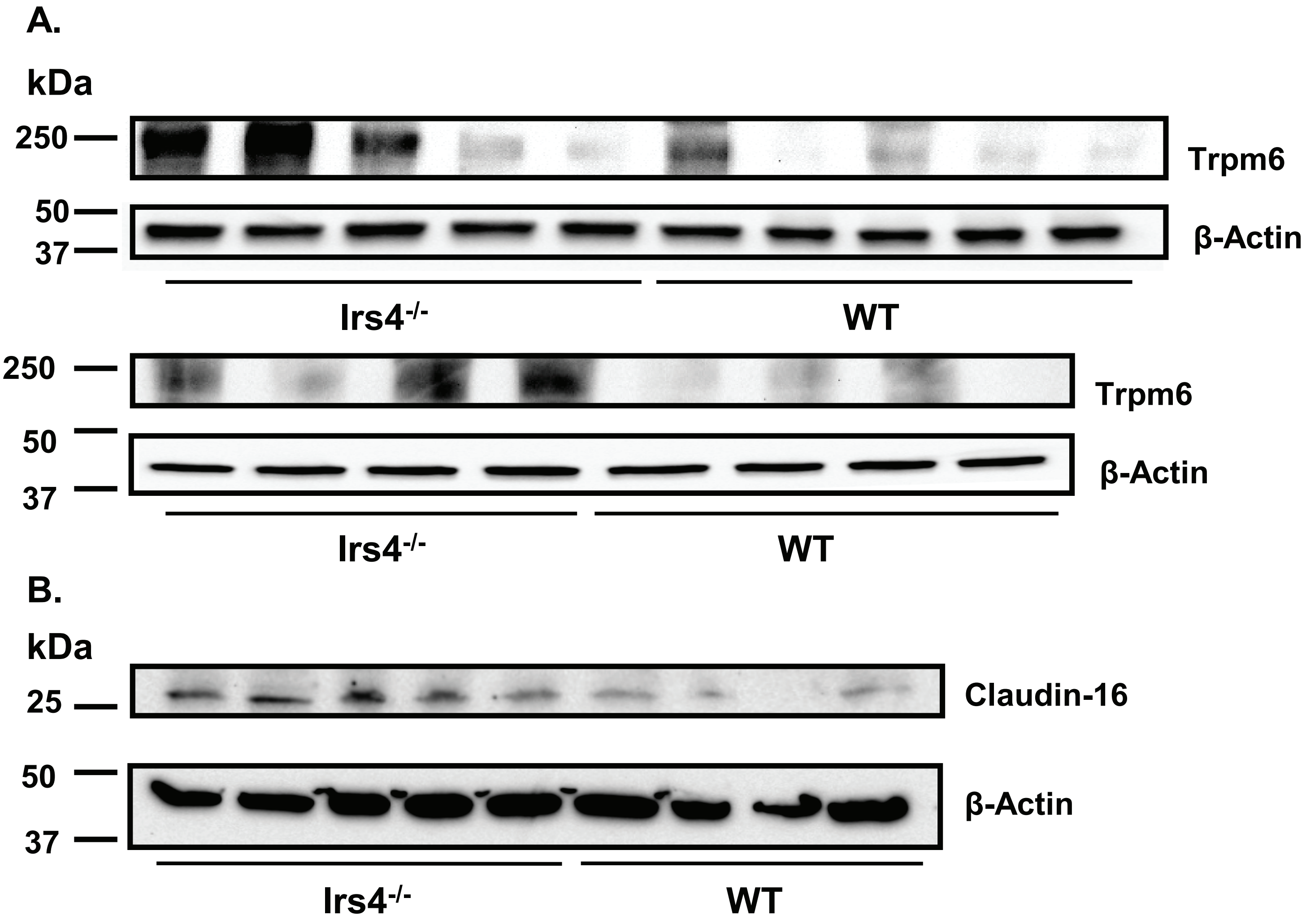
